# Quorum sensing regulation in *Erwinia carotovora* affects development of *Drosophila melanogaster* infected larvae

**DOI:** 10.1101/2019.12.13.876318

**Authors:** Filipe J. D. Vieira, Pol Nadal-Jimenez, Luis Teixeira, Karina B. Xavier

**Affiliations:** Instituto Gulbenkian de Ciência, Oeiras, Portugal; Faculdade de Medicina da Universidade de Lisboa, Lisboa, Portugal

**Author notes:** Address correspondence to: Karina B. Xavier. University of Liverpool, Institute of Integrative Biology, Liverpool, UK.

## Abstract

Multi-host bacteria must rapidly adapt to drastic environmental changes, relying on integration of multiple stimuli for an optimal genetic response. *Erwinia spp.* are phytopathogens that cause soft-rot disease in plants. *Erwinia carotovora Ecc15* is used as a model for bacterial oral-route infection in *Drosophila melanogaster* as it harbors a gene, the *Erwinia* virulence factor (Evf), which has been previously shown to be a major determinant for infection of *D. melanogaster* gut. However, the factors involved in regulation of *evf* expression are poorly understood. We investigated whether *evf* could be controlled by quorum sensing since, in the *Erwinia* genus, quorum sensing regulates pectolytic enzymes, the major virulence factors needed to infect plants. Here, we show that transcription of *evf* is positively regulated by quorum sensing in *Ecc15 via* the acyl-homoserine lactone (AHL) signal synthase ExpI, and the AHL receptors ExpR1 and ExpR2. Moreover, we demonstrate that the GacS/A two-component system is partially required for *evf* expression. We also show that the load of *Ecc15* in the gut depends upon the quorum sensing-mediated regulation of *evf*. Furthermore, we demonstrate that larvae infected with *Ecc15* suffer a developmental delay as a direct consequence of the regulation of *evf via* quorum sensing. Overall, our results show that *Ecc15* relies on quorum sensing to control production of both pectolytic enzymes and Evf. This regulation influences the interaction of *Ecc15* with its two known hosts, indicating that quorum sensing and GacS/A signaling systems may impact bacterial dissemination *via* insect vectors that feed on rotting plants.

**Significance:** Integration of genetic networks allows bacteria to rapidly adapt to changing environments. This is particularly important in bacteria that interact with multiple hosts. *Erwinia carotovora Ecc15* is a plant pathogen that uses *Drosophila melanogaster* as a vector. To interact with these two hosts, *Ecc15* uses two different sets of virulence factors: plant cell wall-degrading enzymes to infect plants and the *Erwinia* virulence factor (*evf*) to infect *Drosophila*. Our work shows that, despite the virulence factors being different, both are regulated by homoserine lactone quorum sensing and the two component GacS/A system. Moreover, we show that these pathways are essential for *Ecc15* loads in the gut of *Drosophila* and that this interaction carries a cost to the vector in the form of a developmental delay. Our findings provide evidence for the importance of quorum sensing regulation in the establishment of multi-host interactions.

## Introduction

Insects play an important role in the dissemination of microorganisms that cause both human and plant diseases. This dissemination may be an active process whereby microbes develop strategies to interact with insects and use them as vectors (1, 2). To do so, bacteria must have the ability to persist within the host (either lifelong or transiently), evading or resisting its immune system in order to abrogate their elimination (3, 4). The host vector will respond with a battery of innate defenses, such as production of antimicrobial peptides and reactive oxygen species as well as behavioral strategies (e.g. avoidance), and physiological responses (e.g. increased peristalsis) (5–9). The successful establishment of these interactions, from the bacterial perspective, ultimately depends on maximizing the fitness of the microorganism and minimizing the impact on the fitness of the vector host (1). Phytopathogenic bacteria such as *Phytoplasma sp., Xylella fastidiosa*, *Pantoea stewartii* (formerly *Erwinia stewartii*), or *Erwinia carotovora* (also known as *Pectobacterium carotovorum*), are among those known to establish close associations with insects and to rely on these hosts as vectors, presumably to facilitate rapid dissemination among plants (10–13). Thus, understanding the molecular mechanisms governing the establishment of these interactions is crucial to prevent insect-borne diseases.

Bacteria from the *Erwinia* genus produce pectolytic enzymes that degrade plant tissue, causing soft root-disease (14). These bacteria survive poorly in soil, overwinter in decaying plant material (14), and use insects, including *Drosophila* species (12, 15) as vectors. Specifically, the non-lethal interaction between the phytopathogen *Erwinia carotovora* (strain *Ecc15*) and *Drosophila melanogaster* has been used as a model to study bacteria-host interactions. Oral infections with *Ecc15* lead to a transient systemic induction of the immune system in *D. melanogaster* and consequent production of antimicrobial peptides (7, 16). These responses are strain-specific and highly dependent on the expression levels of the *Erwinia* virulence factor gene (*evf*) (17), which promotes bacterial infection of the *Drosophila* gut (18). Additionally, expression of *evf* requires the transcriptional regulator Hor (17), but the signals required for the activation of this regulator remain unknown.

Quorum sensing has recently been shown to be important in the regulation of bacterial traits that affect the persistence and/or virulence of bacteria in insects (19–22). Many bacteria use quorum sensing to regulate gene expression as a function of population density (23, 24). This cell-cell signaling mechanism relies on the production, secretion, and response to extracellular signaling molecules called autoinducers (24–26). Bacteria from the *Erwinia* genus produce a mixture of plant cell wall-degrading enzymes (PCWDE), which are the major virulence factors used to degrade plant tissues and potentiate bacterial invasion of the plant host (27–30). In these bacteria, expression of these PCWDE is tightly regulated by two main signaling pathways: the acyl-homoserine lactone (AHL) quorum sensing system, and the GacS/A two-component system (31–34). Typically, the AHL quorum sensing system present in *Erwinia spp.* includes the AHL synthase ExpI (35), and two AHL receptors, ExpR1 and ExpR2 (36), which are homologues to the canonical LuxI/R quorum sensing system first identified in *Vibrio fischeri* (37–39). The GacS/A two-component system is also activated at high cell density, and, like the AHL quorum sensing system, regulates virulence in many Gram-negative pathogenic bacteria (40–45). Given the importance of these two signal transduction pathways for the expression of the major plant virulence factors in *Erwinia spp.,* we investigated whether quorum sensing and the GacS/A system also regulate *evf* expression in *Ecc15*. Additionally, we tested whether these signaling pathways are important for *Ecc15* infection, and determined the consequences of this interaction for the insect host. Our results show that PCWDE and *evf* expression in *Ecc15*, which are required for the interactions with plants and insects, respectively, are both regulated by the same quorum sensing signalling pathway. Moreover, we demonstrate that *evf* expression has a negative effect on the insect host as it leads to a developmental delay in larvae infected with *Ecc15*.

## Results

### The expression of *evf* is regulated by both AHL-dependent quorum sensing and the GAC system

We first investigated whether activation of the production of PCWDE in *Ecc15* requires both the AHL quorum sensing system and the GacS/A two-component system (GAC), as occurs in other members of the *Erwinia* (or *Pectobacterium*) genus (32, 35, 46). We constructed deletion mutants of *expI* and *gacA,* the genes encoding homologues of the AHL-synthase and the response regulator of the GAC system, respectively. We determined whether any of these mutations cause a growth defect in *Ecc15*, and observed no difference in growth compared to the WT strain (Fig. S1). We then measured pectate lyase activity in supernatants of cultures from *Ecc15* WT, *expI* or *gacA* mutants, as this is one of the PCWDE typically secreted by *Erwinia spp.*. As shown in Fig. 1a (and replicate experiments in Fig. S2), both the *expI* and the *gacA* mutants exhibit pronounced reductions in pectate lyase activity when compared to the WT (TukeyHSD test, *p*<0.001, Fig. S2C). Addition of a mixture of exogenous 3-oxo-C6-HSL and 3-oxo-C8-HSL, the major AHLs produced by *Erwinia carotovora* (46), to an *expI* mutant culture was sufficient to restore production of this PCWDE to higher levels than the WT (Fig. 1A, TukeyHSD test *p*<0.001, Fig. S2C). In addition, both the *expI and gacA* mutants are impaired in virulence to the plant host, which we tested by measuring the mass of macerated tissue in potato tubers inoculated with these genotypes (Fig. 1B, TukeyHSD test *p*<0.001, Fig. S2F). In contrast, the *evf* mutant shows no significant difference in maceration with respect to the WT (Fig. 1B and Fig. S2D-F). Altogether, these results show that production of pectate lyase, as well as plant host-virulence, are regulated by both the AHL and GAC systems in *Ecc15*, as occurs in other *Erwinia spp.,* where *expI* and *gacA* mutants have been shown to be avirulent (34, 47, 48). Moreover, we show that *evf* is not necessary for plant infection (Fig. 1B and Fig. S2D-F).

**Fig. 1.**
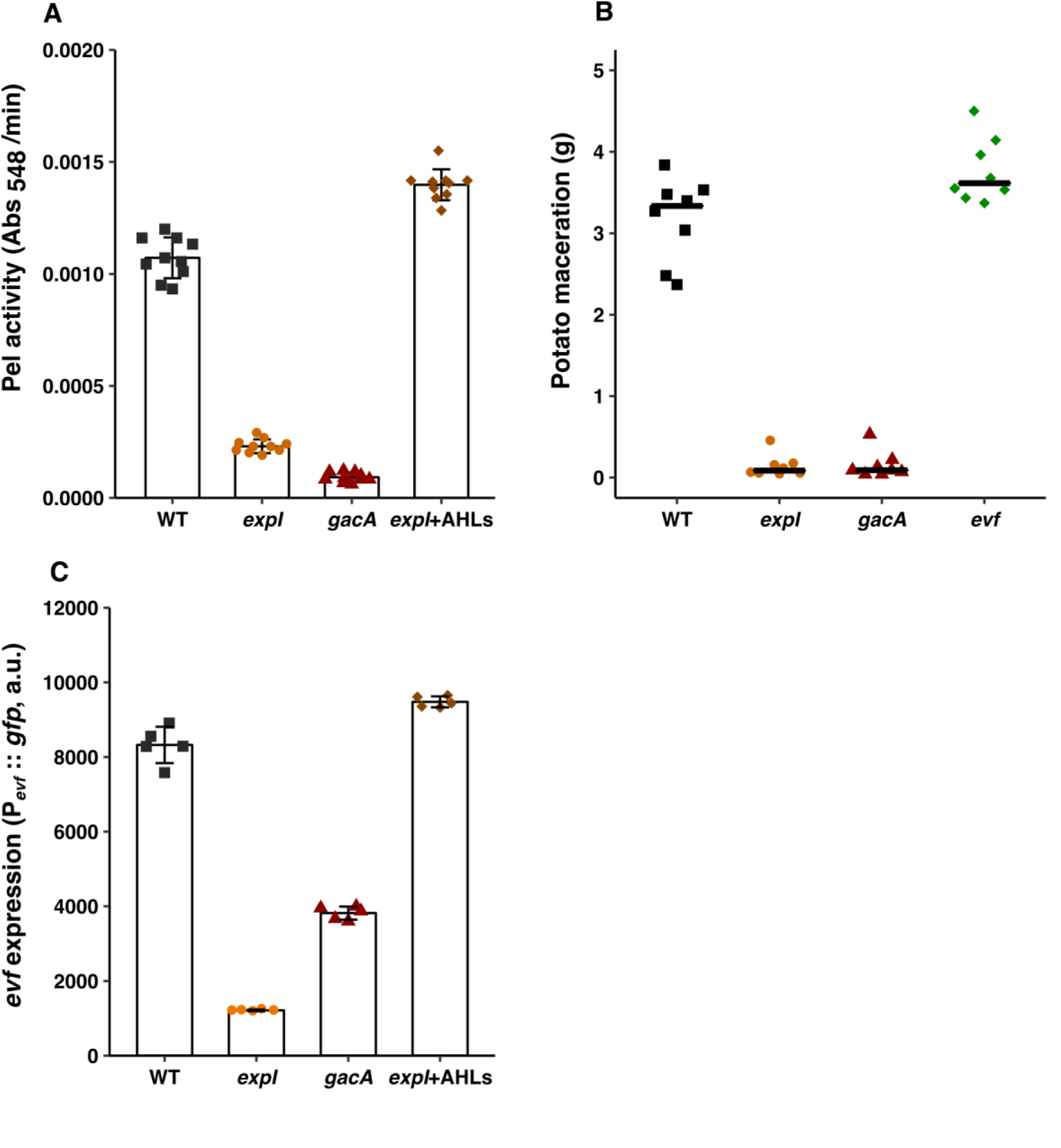
Production of pectate lyase and expression of *evf* is dependent on both quorum sensing and the GAC system. **(A)** Pectate lyase activity in cell-free supernatants of WT *Ecc15*, *expI* and *gacA* mutants at 6 hours of growth in LB + 0.4%PGA. n=10 **(B)** Potato maceration quantification (grams) in potatoes infected with WT *Ecc15*, *expI*, *gacA* and *evf* mutants, 48 hours post-infection. n=8 **(C)** Pevf::*gfp* expression in WT *Ecc15*, *expI* and *gacA* mutants at 6 hours of growth in LB + Spec. n=5 Growth curves of the strains used are shown in Fig.S1. Complementation with AHLs was performed with a mixture of 1uM 3-oxo-C6-HSL and 3-oxo-C8-HSL. Error bars represent standard deviation of the mean. For each panel a representative experiment from three independent experiments is shown (other two experiment are shown in Fig. S2). Statistical analysis taking the data of all the three experiments is shown in Fig. S2.

To investigate whether *evf* expression is also regulated by these two systems, we analyzed the expression of a transcriptional reporter consisting of a Green Fluorescent Protein (GFP) fused to the promoter of *evf* (P*_evf_*::*gfp*) in mutants of either AHL quorum sensing or GAC signaling systems. We observed that the expression of the P*_evf_*::*gfp* is reduced in the *expI* mutant when compared to the WT (TukeyHSD test, *p*<0.001), and that this expression can be restored if exogenous AHLs are supplied to the culture (Fig. 1C, Fig. S2G-I). In the *gacA* mutant, expression of the *evf* promoter is also reduced compared to the WT, but not as much as in the *expI* mutant (Fig. 1C, TukeyHSD test *p*<0.001, Fig. S2G-I). Since it was previously shown that mutants in the GAC system produce less AHLs (34), we asked if the difference observed between the WT and the *gacA* mutant could be solely explained by the lower levels of AHLs produced by the latter. However, addition of exogenous AHLs to the cultures of a *gacA* mutant did not restore the levels of P*_evf_*::*gfp* expression to WT levels (Fig. S3). Therefore, we conclude that the *gacA* phenotype regarding *evf* expression is mostly independent of AHLs. Overall, these results show that full activation of both *evf* expression and PCWDE activity is dependent on quorum sensing regulation *via* AHLs, and, to a lesser extent, on activation of the GAC system.

In the absence of AHLs, the AHL receptors ExpR1 and ExpR2 lead to repression of virulence traits such as PCWDE (35, 49). These receptors are DNA binding proteins that act as transcriptional activators of *rsmA,* which encodes a global repressor of quorum sensing-regulated genes in *Erwinia spp.* (36, 49, 50). Upon AHL binding, these receptors lose their ability to bind DNA, resulting in decreased expression of *rsmA* and, consequently, increased expression of virulence traits (51, 52). To determine whether ExpR1 and ExpR2 also mediate AHL-dependent regulation of *evf* expression, we constructed deletions of these two genes in the *expI* background. We measured expression of the P*_evf_*::*gfp* reporter in this *expI expR1 expR2* triple mutant, with or without exogenous AHLs. Because AHLs block activation of RsmA *via* ExpR1 and ExpR2 (51, 52), deletion of *expR1 and expR2* in the *expI* background is expected to result in the de-repression of *evf*. Consistent with this prediction, P*_evf_*::*gfp* expression is higher in the *expI expR1 expR2* than in the *expI* single mutant (Fig. 2A, TukeyHSD test *p*<0.001, Fig. S4A-C). However, the expression levels of P*_evf_*::*gfp* are lower in the *expI expR1 expR2* than those of the WT (Fig. 2A, TukeyHSD test, *p*<0.001, Fig. S4A-C). The fact that deletion of these two receptors in the *expI* background is not sufficient to fully restore expression of *evf* to WT levels indicates that additional regulators control the expression of *evf*. Nonetheless, while addition of exogenous AHLs to a culture of an *expI* mutant increases P_evf_::*gfp* expression, it remains unaltered in the triple *expI expR1 expR2* mutant (Fig. 2A, TukeyHSD test *p*=1, Fig. S4A-C). Therefore, AHL-dependent regulation of *evf* expression is mediated by *expR1* and *expR2,* as is also the case for the regulation of PCWDE in other *Erwinia spp.* (34, 36, 49).

**Fig. 2.**
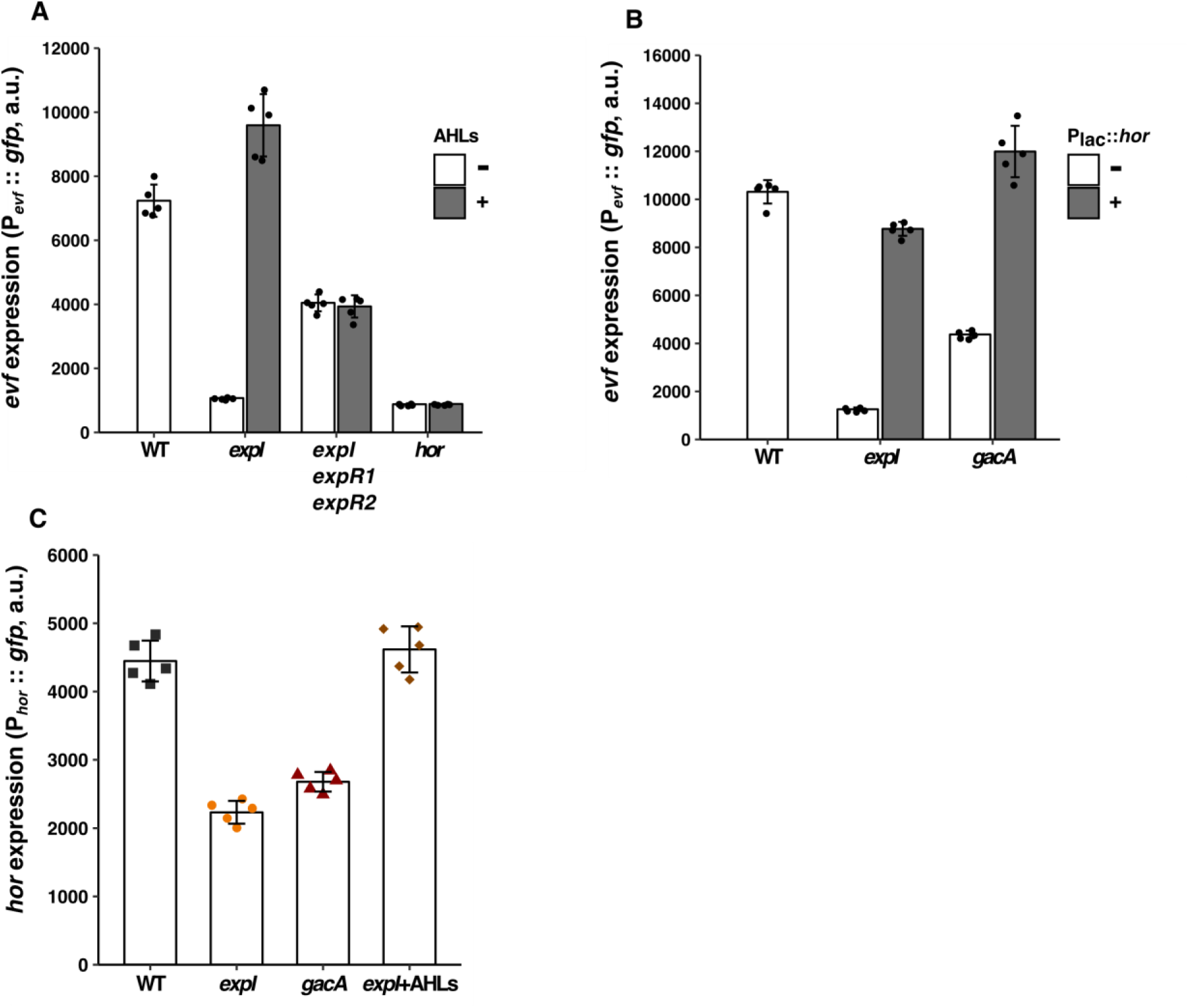
*evf* regulation by quorum sensing is dependent on ExpR receptors and *hor*. **(A)** P*_evf_*::*gfp* expression without (white bars) or with (grey bars) addition of exogenous AHLs in *Ecc15*, *expI*, *expI expR1 expR2* and *hor* mutants at 6 hours of growth in LB + Spec. n=5 **(B)** P*_evf_*::*gfp* expression in *Ecc15 expI* and *gacA* mutants containing a plasmid with the P*_evf_*::*gfp* fusion (white bars)or with both P*_lac_*::*hor* and P*_evf_*::*gfp* fusions (grey bars) at 6 hours of growth in LB + Spec. n=5 **(C)** P*_hor_*::*gfp* expression in WT *Ecc15*, *expI* and *gacA* mutants at 6 hours of growth in LB + Spec. n=5 Complementation with AHLs was performed with a mixture of 1µM 3-oxo-C6-HSL and 3-oxo-C8-HSL. Error bars represent standard deviation of the mean. For each panel a representative experiment from three independent experiments is shown (other two experiment are shown in Fig. S4). Statistical analysis taking the data of all the three experiments is shown in Fig. S4.

### Regulation of *evf* by AHL quorum sensing is mediated by *hor*

It was previously shown that Hor, a global regulator of diverse physiological processes in many animal and plant bacterial pathogens (53), is a positive regulator of *evf* (17) and that, as in other *Erwinia spp.*, *hor* is regulated by quorum sensing (54). Therefore, we asked if AHL-dependent regulation of *evf* is *via hor*. We analyzed the expression of the P*_evf_*::*gfp* reporter in a *hor* mutant, and found that it is lower than in the WT, and as low as in the *expI* mutant (Fig. 2A). Moreover, we observed that addition of exogenous AHLs to a *hor* mutant does not restore the expression of *evf* (Fig. 2A, TukeyHSD test *p*=1, Fig. S4A-C). We next cloned the *hor* gene under the control of a *lac* promoter in the plasmid containing the P*_evf_*::*gfp* fusion, and measured *evf* expression levels in the *expI* and *gacA* mutants expressing or not the *hor* gene. We observed that expression of *hor* in either the *expI* or the *gacA* mutants restores *evf* expression to levels similar to those of the WT (Fig. 2B, TukeyHSD test *p*<0.001, Fig. S4D-F). Therefore, regulation of *evf* is mediated by both the AHL and the GAC systems and occurs via *hor*. Next, we asked whether these systems regulate *hor* itself by analyzing the expression of a *hor* promoter fusion (P*_hor_*::*gfp*) in *expI* and *gacA* mutants. As for the *evf* reporter, we observed that, P*_hor_*::*gfp* expression is lower in an *expI* mutant when compared to the WT (Fig. 2C, TukeyHSD test *p*<0.001, Fig. S4G-I). Moreover, this expression can be complemented to WT levels by the addition of exogenous AHLs to the growth medium of the *expI* mutant (Fig. 2C, TukeyHSD test *p*=0.08, Fig. S4G-I). These data demonstrate that *hor* expression is regulated by AHLs and is necessary for the increase of *evf* expression mediated by AHLs.

### Infection by *Ecc15* causes a developmental delay in *D. melanogaster* larvae dependent on quorum sensing and GAC regulation of *evf* expression

It is known that Evf promotes infection in the *D. melanogaster* gut (18, 19). To examine the effects of down-regulation of *evf* on quorum sensing and GAC mutants we measured *Ecc15* loads upon oral infection. We inoculated *Ecc15* WT, *evf*, *expI* or *gacA* into *D. melanogaster* L3 stage larvae, and assessed the dynamics of bacterial loads by counting the number of colony forming units (CFU) of *Ecc15* over time. As previously reported, *Ecc15* infection is transient and larvae are able to clear it after 24 hours (Fig. 3 and (18)). Additionally, we observed that the rate of elimination of the bacteria from the larval gut is not significantly different between the WT and the *evf*, *gacA*, and *expI* mutants (Fig3, lmm, Chi-square test *p*=0.27). However, we also observed that *Ecc15* WT loads were approximately ten times higher compared to the loads of the *evf* mutant when considering the entire infection period (Fig. 3, TukeyHSD test *p*<0.001, Fig. S5), confirming that *evf* is required for optimal infection of the larval gut by *Ecc15*. Importantly, a similar trend was observed when comparing the WT to either of the two mutants impaired in *evf* expression: *gacA* or *expI* (Fig. 3. TukeyHSD test *p*<0.001, Fig. S5), revealing the importance of quorum sensing-regulation and the GAC system in the infection process. Taken together, our data show that *evf* provides *Ecc15* with the ability to reach high loads in the insect gut, but does not increase its capacity to survive inside it.

**Fig. 3.**
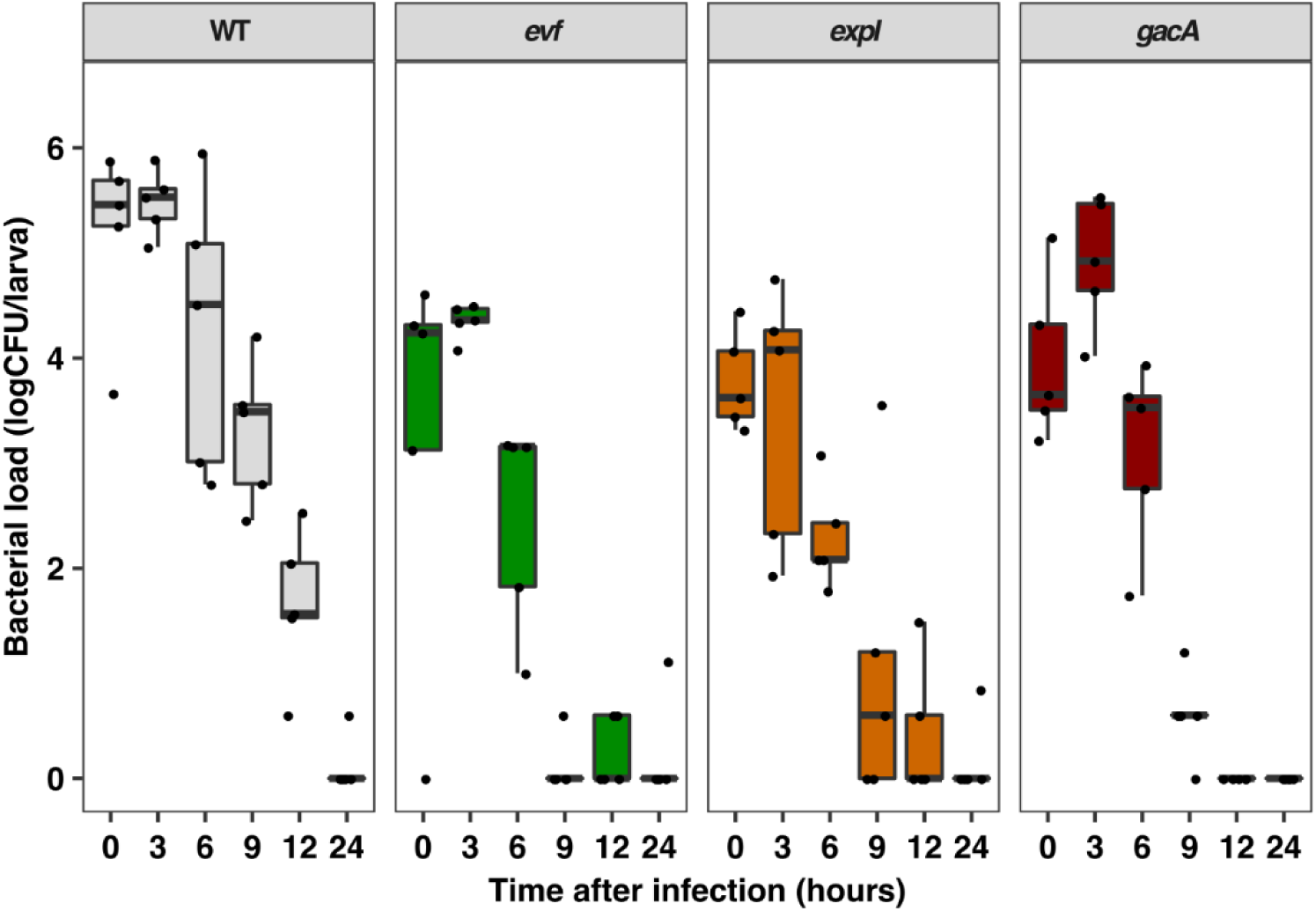
*Ecc15* loads are higher in *D. melanogaster* larvae orally infected with WT than with mutants impaired in *evf* expression. *D. melanogaster* L3 stage larvae were infected with WT *Ecc15*, *evf, expI* and *gacA* mutants for 30 min and then transferred to fresh media. Following the infection period Colony Forming Units (CFUs) of *Ecc15* were measured at the specified time points. Each dot represents CFUs of one single larvae (5 larvae per time point). 0 hours after infection correspond to 30 min of confined exposure to 200µl of an OD_600_=200. Representative experiment from three independent experiments (other two experiment are shown in Fig. S5). Statistical analysis of the comparison of the entire infection period for each condition tested using the data of all the three experiments is shown in Fig. S5.

Next, we asked if infection of *D. melanogaster* larvae by *Ecc15* has an effect on larval development. To investigate this possibility, we infected *D. melanogaster* L3 stage larvae orally with *Ecc15* WT or an *evf* mutant and followed their development over time. We found that infection by WT *Ecc15* delays *D. melanogaster* larvae passage to pupal stage an average of 49 hours, when compared to non-infected larvae (Fig. 4A and FigS6, TukeyHSD test *p*<0.001, Fig. 4B). Moreover, we show that this strong delay is *evf-* dependent, since larvae exposed to an *evf* mutant only show a delay of 8 hours when compared to non-infected larvae (TukeyHSD test, *p*<0.001, Fig4B). We then asked if the mutants in the quorum sensing pathway and GAC system, which have low expression of *evf*, would show a similar phenotype. We observed that larvae exposed to the *expI* mutant, which has very low expression of *evf*, also show only a 4 hour delay with respect to non-infected larvae, similar to the *evf* mutant (TukeyHSD test, *p*<0.001, Fig4B). Interestingly, larvae infected with the *gacA* mutant, which has intermediate levels of *evf* expression, show an intermediate developmental delay, taking an average of 26 hours longer than non-infected larvae to reach the pupal stage (TukeyHSD test, *p*<0.001, Fig4B). Since the developmental delay correlated with the levels of *evf* expression in the strains tested, we next examined whether constitutive overexpression of *evf* would exacerbate the phenotype. We observed that larvae infected with a WT *Ecc15* overexpressing *evf* died before reaching the pupal stage (Fig. 4C-D). These results show that *Ecc15* has a negative impact on larval development and this effect requires both *evf* and the quorum sensing and GAC regulatory systems.

**Fig. 4.**
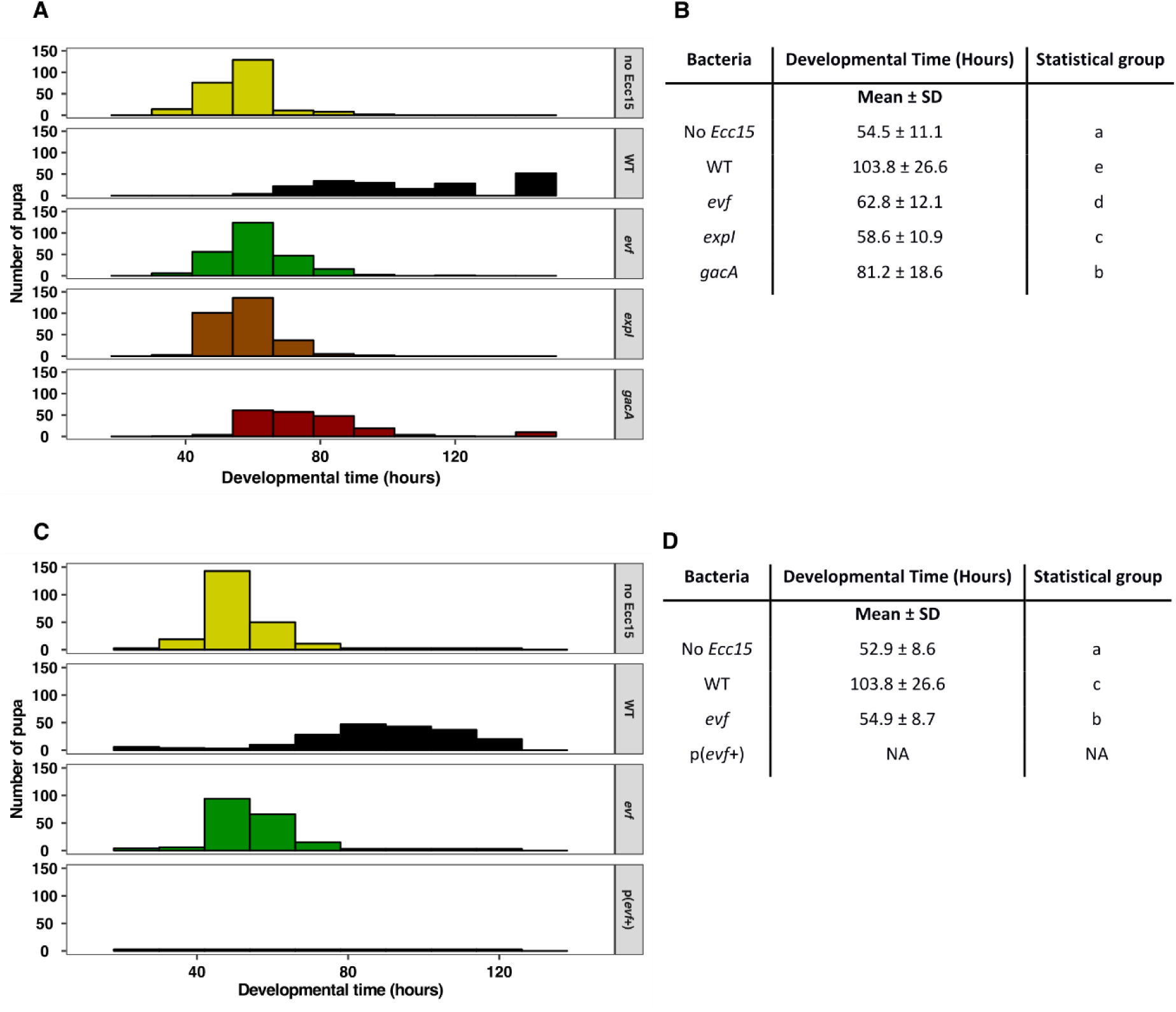
*Ecc15* causes a developmental delay in *D. melanogaster* larvae that is dependent on *evf*, quorum sensing and the GAC system. L3 stage *Drosophila* larvae pupariation time after exposure to **(A)** WT *Ecc15*, *evf*, *expI* and *gacA* mutants or **(C)** WT *Ecc15* overexpressing Evf, compared with non-infected larvae. **(B)** and **(D)** Average developmental time in hours with standard deviation. Representative experiment from three independent experiments (other two experiment are shown in Fig. S6). Statistical groups shown in (B) and (D) were determined using a linear mixed effect model taking in consideration the data from the three experiments. A Tukey HSD test was applied for multiple comparisons using the estimates obtain from the model.

## DISCUSSION

*Erwinia spp.* are phytopathogenic bacteria thought to depend on insects to spread among plant hosts (1, 12, 13). To interact with both plants and insects, *Ecc15* relies on different traits that seem to be specific for the interaction with each host. In this bacterium, PCWDE are the major virulence factors required for plant infection (40) and Evf is required to infect *D. melanogaster,* but not necessary to infect potato tubers (Fig. 1B and (16, 17)). It was not known whether *Ecc15*, which relies on multiple hosts for survival, regulates host-specific traits using the same or different signal transduction networks. Here we showed that the AHL-dependent ExpI/ExpR system, which regulates plant virulence factors (33, 35, 36, 49) is also essential for the expression of the insect virulence factor *evf*, suggesting that the signal transduction networks regulating traits required across hosts are the same. An *expI* mutant had lower levels of *evf* expression than the WT which could be restored by addition of exogenous AHLs to the growth medium. We also demonstrated that the GAC system, that is thought to respond to the physiological state of the cell (42) and is involved in regulation of plant virulence factors (41, 55) is also necessary for full expression of *evf*. Additionally, we showed that regulation by these two networks occurs through *hor*, a conserved transcriptional regulator of the SlyA family (56), previously found to be regulated by quorum sensing in another *E. carotovora* strain (54). ExpR1 and ExpR2 AHL receptors function as activators of *rsmA*, the global repressor of the AHL-regulon; therefore, we expected the *expI expR1 expR2* mutant to have the same levels of *evf* expression as the *expI* mutant supplemented with AHLs. However, we found that the *expI expR1 expR2* mutant has lower levels of *evf* expression than both the *expI* supplemented with AHLs and the WT. Moreover, we showed that complementation of the *expI expR1 expR2* mutant with AHLs does not change the level of *evf* expression. These results show that *expR1* and *expR2* are required for *Ecc15’s* response to AHLs, but also indicate that an additional AHL-independent regulator, is playing a role in the regulation of *evf* in this bacterium. One possibility is that *Ecc15* has additional orphan *luxR* genes, DNA binding proteins homologous to LuxR that lack a cognate AHLs synthase. These orphan genes are divided in two categories, those that have both a LuxR DNA and an AHL binding domain, such as ExpR2, and those that have only the typical LuxR DNA binding domain (57), such as *vqsR* in *Pseudomonas aeruginosa.* In this bacterium, in response to an unknown signal, *vqsR* has been found to downregulate expression of virulence through binding to the promoter region of the quorum sensing receptor *qscR,* inhibiting its expression without responding to AHLs (58). Because addition of exogenous AHLs to the *expI expR1 expR2* mutant does not change the level of *evf* expression, this unknown regulator is more likely to lie within the second category of orphan LuxR receptors. Our data also suggests that this unknown regulator could be repressed by *rsmA,* since the *expI* mutant shows lower levels of *evf* expression than *expI expR1 expR2*. In *Erwinia spp.* another layer of regulation required for PCWDE expression is the detection of external environmental signals like pectin, a component of the plant cell wall (34, 55, 59, 60, 35, 51, 47, 48). In the absence of plant signals, transcription of PCWDE is repressed. Unlike in the regulation of PCWDE in *Erwinia spp.*, in our experimental setting we have no evidence for the need of a host signal since we can detect *evf* expression in cells grown in LB without the need for other signals. However, this does not exclude the possibility that environmental signals, perhaps related to insect derived compounds, have a role in the overall levels of *evf* expression.

It has been hypothesized that *evf* was horizontally acquired by *Ecc15* and a few other *Erwinia spp*. As these phytopathogens often use insects as vectors, one hypothesis for the selective benefit of acquiring *evf* is that this gene might be important to favor bacterial transmission by strengthening the interaction of *Ecc15* with *Drosophila*. This hypothesis is supported by our results showing that *evf* allows *Ecc15* to have higher loads at the initial stage of *Drosophila* larval infection. However, the rate of *Ecc15* elimination post-infection was the same in WT and an *evf* mutant. This suggests that *evf* is promoting transmission of *Ecc15* by increasing the overall number of bacteria that reach the gut. Moreover, we show here that larvae infected with *Ecc15* are developmentally impaired when compared to non-infected larvae, and this developmental delay is dependent on *evf.* These results are in agreement with previous reports showing that larvae infected with WT *Ecc15* were smaller due to inhibition of the larval proteolytic activity promoted by *Drosophila*-associated *Lactobacillus* species (61). Additionally, infection of *Drosophila* adults and larvae with WT *Ecc15* causes cell damage, which induces epithelial cell death, leading to activation of immunity, stem cell regeneration programs and differentiation/modification of the cellular structure of the gut, essential for its repair (7, 16, 62). These studies, together with our results, show that *evf* expression in *Ecc15* has an overall deleterious effect on the host, and thus acquisition of *evf,* which enables higher host loads and is presumably beneficial for bacterial transmission, seems to have resulted in a tradeoff for host fitness.

Due to a lack of genetic information, tracing the evolutionary history of this protein is challenging. It was previously reported that, besides *Ecc15*, *evf* was only identified in strain *Ecc1488* (16, 17). By comparing the amino acid sequence of Evf to recent genome databases, we found only a few more candidate ortholog proteins with amino acid sequence identity higher than 60% (Table S3). The highest sequence similarities found, besides those of other *Erwinia spp.*, corresponded to proteins from *Cedecea neteri, Enterobacter AG1, Rahnella sp., Klebsiella aerogenes* and *Escherichia coli* (Table S3). *K. aerogenes and E. coli* are ubiquitous bacterial species that can colonize the gut of different animals, particularly mammals, but also insects (63–65). Similarly to *Erwinia spp.*, *Rahnella sp*. and *C. neteri* are bacterial species often isolated from plants that also establish gut associations with insects (64, 66, 67). Enterobacter AG1 is a bacterial species isolated from the gut of mosquitos that has been shown to decrease the ability of *Plasmodium falciparum* to colonize the gut (68, 69). Since the structural fold of Evf is unique (70) and that protein structure is more conserved than sequence identity (71), we predicted the secondary structures of these ORFs using phyre2 (72). We found that the predicted secondary structure of all five ORFs is identical to Evf (Table S3). Importantly, the cysteine residue (position 209), which in *Ecc15* Evf is palmitoylated, a post-translational modification essential for its function (70), is conserved in all the five ORFs. Interestingly, *evf*-like genes with low amino acid sequence identity (lower than 40%), but with a predicted secondary structure highly similar to that of the Evf (72), can be found in other bacteria such as *Vibrio sp.* or the major insect pathogen *Photorhabdus luminescens* ((18) Locus PLU2433). *P. luminescens* colonizes the gut of *Heterorhabditis bacteriophora,* an insect-preying nematode (73, 74). The nematode enters through the insect’s respiratory and/or digestive tract and regurgitates the bacteria into its hemolymph. Once in the hemolymph, *Photorhabdus* produces a battery of toxins that kills the insect allowing the nematode to feed on the corpse, favoring *Photorhabdus* recolonization (75–77). *Photorhabdus* possesses several genes possibly involved in the establishment of the interaction with the host, many of which are regulated by quorum sensing (78, 79). Thus, it is possible that the Evf ortholog from *Photorhabdus* is involved in the mechanisms required for colonization of the nematode, or in the pathogenicity towards the insect. Our results indicate that Evf orthologs can be found in bacteria with apparently different lifestyles. However, all of these bacteria encounter multiple hosts mainly through the gut, including insects, and undergo rapid environmental changes. It is possible that Evf has a conserved role in host transition mainly through insect colonization or pathogenesis.

Quorum sensing regulation is associated with tight control of density dependent activation of genes encoding functions that are often essential for the establishment of host-microbe interactions (26). For instance, in the interaction between the squid *Euprymna scolopes* and *V. fischeri,* mutants in the quorum sensing system are less efficient in persisting in the light organ, being outcompeted by other strains (80, 81). Here we show that in *Ecc15*, besides regulating PCWDE in plant infections, employs quorum sensing for the *evf*-mediated increased bacterial loads in *Drosophila* larvae. Our study also demonstrates that the quorum sensing and GAC regulatory pathways have a strong effect in the Evf-mediated developmental delay caused by *Ecc15.* Moreover, overexpression of *evf* leads to a complete developmental arrest of larvae, eventually killing them. Therefore, one possible benefit of having *evf* expression under the control of these networks might be to minimize the detrimental effect that the *evf*-dependent infection has on the insect host while still enabling a transient infection. On the other hand, insects are attracted to rotten plant tissue, and if *evf* is important for promoting the interaction of *Ecc15* with its insect vector (*Drosophila)*, synchronization of the expression of *evf* and the PCWDE might have been selected as advantageous for bacterial dissemination. This phenomenon, called predictive behavior, is particularly common in symbiotic relationships where the microbe often experiences a predictable series of cyclic environments (82). In mammalian hosts, a very predictable change when transitioning from the outside environment to the oral cavity is the immediate increase in temperature followed by a decrease in oxygen. This phenomenon has been described for *E. coli* gut colonization where, coupled to an increase in temperature, downregulation of genes related to aerobic respiration is observed (83). In the case of *Ecc15* it is possible that control of PCWDE and *evf* expression is intertwined so that following colonization of the plant, *evf* expression is triggered, anticipating the appearance of the insect vector which is attracted to rotten plant tissue, and thus maximizing the probability of establishing the interaction with this host vector.

Our results show that, in *Ecc15*, the regulatory networks responding to self-produced quorum sensing signals and physiological cues sensed by the GAC system are used to control expression of traits required to infect different hosts. Thus, the signal transduction mechanisms are the same even though the functions involved in the interactions with each plant or insect host are largely different. Therefore, our findings reinforce the central role of quorum sensing in the regulatory circuitry controlling the array of traits used by bacteria to interact with diverse hosts.

## Materials and Methods

### Bacterial strains, plasmids, and culture conditions

The strains and plasmids used in this study are listed in Table S1 of the supplementary material. All bacterial strains used are derived from wild type (WT) *Ecc15* strain (7). *Ecc15* and mutants were grown at 30°C with aeration in Luria-Bertani medium (LB). When specified, medium was supplemented with 0.4% polygalacturonic acid (PGA; Sigma P3850), to induce the expression of PCWDEs. *E. coli* DH5α was used for cloning procedures and was grown at 37°C with aeration in LB. When required, antibiotics were used at the following concentrations (mg liter^−1^): ampicillin (Amp), 100; kanamycin (Kan), 50; spectinomycin (Spec), 50; chloramphenicol (Cm), 25. To assess bacterial growth, optical density at 600 nm (OD_600_) was determined in a Thermo Spectronic Helios delta spectrophotometer.

### Genetic and molecular techniques

All primer sequences used in this study are listed in Table S2 in supplemental material. *P. carotovorum Ecc15* deletion mutants listed in Table S1 were constructed by chromosomal gene replacement with an antibiotic marker using the λ-Red recombinase system (84). Plasmid pLIPS, able to replicate in *Ecc15* and carrying the arabinose-inducible λ-Red recombinase system was used (34). Briefly, the DNA region of the target gene, including approximately 500 bp upstream and downstream from the gene, was amplified by PCR and cloned into pUC18 (85) using restriction enzymes. These constructs, containing the target gene and its flanking regions, were divergently amplified by PCR, to introduce a *Xho*I restriction site in the 5′ and 3′ regions and to remove the native coding sequence of the target gene. The kanamycin cassette from pkD4 was amplified with primers also containing the *Xho*I restriction site. The fragment containing the kanamycin cassette was then digested with *Xho*I and was introduced into the *Xho*I-digested PCR fragment carrying the flanking regions of the target gene. The final construct, containing the kanamycin cassette flanked by the upstream and downstream regions of the target gene was then amplified by PCR, and approximately 2 micrograms of DNA were electroporated into the parental strain (FDV31) expressing the λ-Red recombinase system from pLIPS, to favour recombination. To construct the plasmid carrying the promoter *evf* fused to GFP (pFDV54), a fragment of 503 bp containing the *evf* promoter was amplified from WT *Ecc15* DNA with the primers P1194 and P1195. This fragment was then digested with *Hin*dIII and *Sph*I and ligated to pUC18. GFP was amplified from the pCMW1(86) vector using primer P0576 and P0665. Both the GFP and pUC18-P_evf_ were digested with *Sph*I and *Bam*HI, ligated and 2 µl of the ligation reaction were used to transform Dh5α (pFDV54). The same procedure was used for the P*_hor_*::*gfp* fusion using primers P1351 and P1352 for promoter amplification (493 bp) and primers P1353 and P1354 for GFP amplification. Digestions were made with enzymes *Hin*dIII/*Pst*I and *Pst*I/*Xba*I (pFDV84). For *hor* overexpression, a *Nco*I site was introduced in pOM1-P*_evf_*::*gfp* with primers P1309 and P1310. *hor* was amplified using primers P1311 and Primers 1312 from WT template DNA. Then both the plasmid and the fragment carrying *hor* were digested with *Nco*I and *Sac*I and subsequently ligated (pFDV104).

PCR for cloning purposes was performed using the proofreading Bio-X-ACT (Bioline) enzyme. Other PCRs were performed using Dream Taq polymerase (Fermentas). Digestions were performed with Fast Digest Enzymes (Fermentas), and ligations were performed with T4 DNA ligase (New England Biolabs). All cloning steps were performed in either *E. coli* DH5α or WT *Ecc15*. All mutants and constructs were confirmed by PCR amplification and subsequent Sanger sequencing performed at the Instituto Gulbenkian de Ciência sequencing facility.

### Pectate lyase activity assay

*Ecc15* and mutants were grown overnight in LB with 0.4% PGA, inoculated into fresh media to a starting OD_600_ of 0.05 and incubated at 30°C with aeration. After 6 hours of incubation, aliquots were collected to evaluate growth and to analyse pectate lyase (Pel) activity in cell-free supernatants, using the previously described procedure (55) based on the thiobarbituric acid colorimetric method (87). Each experiment included at least 5 independent cultures per genotype, and was repeated on 3 independent days.

### Plant virulence assay

Plant virulence was analysed by assessing the maceration of potato tubers with the protocol adapted from (34, 88). Potatoes were washed and surface sterilized by soaking for 10 min in 10% bleach, followed by 10 min in 70% ethanol. Overnight cultures in LB broth were washed twice and diluted to an OD600 of 0.05 in phosphate-buffered saline (PBS). Thirty-microliter aliquots were then used to inoculate the previously punctured potatoes. Potato tubers were incubated at 28°C at a relative humidity above 90% for 48 h. After incubation, potatoes were sliced, and macerated tissue was collected and weighed.

### Promoter expression assays

*Ecc15* carrying the different plasmid-borne promoter reporter fusions were grown overnight in LB supplemented with Spectinomycin (LB + Spec), inoculated into fresh medium at a starting OD_600_ of 0.05 and incubated at 30°C with aeration. At the indicated timepoints, aliquots were collected to assess growth and the expression of the reporter fusion. For the analyses of reporter expression, aliquots of the cultures were diluted 1:100 in PBS and expression was measured by flow cytometry (LSRFortessa; BD) and analysed with Flowing Software v 2.5.1, as previously described (55). A minimum of 10,000 green fluorescent protein (GFP)-positive single cells were acquired per sample. Expression of the promoter-*gfp* fusions is reported as the median GFP expression of GFP-positive single cells in arbitrary units. Each experiment included at least 5 independent cultures per genotype, and was repeated on 3 independent days.

### Drosophila Stocks

DrosDel *w^1118^* isogenic stock (*w^1118^ iso*) was used in all experiments (89, 90). Stocks were maintained at 25°C in standard corn meal fly medium composed of 1.1 L water, 45 g molasses, 75 g of sugar, 10 g agar, 70 g cornmeal, 20 g yeast. Food was autoclaved and cooled to 45°C before adding 30 mL of a solution containing 0.2 g of carbendazim (Sigma) and 100 g of methylparaben (Sigma) in 1 L of absolute ethanol. Experiments were performed at 28°C

### Developmental delay and bacterial CFUs assays

Egg laying was performed in cages containing adult flies at a ratio of 3 females to 1 male. To synchronize the embryo stage, flies were initially incubated for 1 hour at 25°C to lay prior fertilized eggs. After this initial incubation, flies were transferred to new cages where eggs were laid for 4 to 6 hours in the presence of standard corn meal fly medium. After this period, eggs were removed and incubated at 25°C for 72 hours to obtain L3-stage larvae. For bacterial infections, third-instar larvae were placed in a 2 ml Eppendorf containing 200 µl of concentrated bacteria pellet (OD_600_ = 200) from an overnight culture and 400 µl of standard corn meal fly medium. Larvae, bacteria and food were then thoroughly mixed with a spoon, the Eppendorf was closed with a foam plug and incubated at room temperature for 30 min. The mix was then transferred to a 25 ml plastic tube containing 7.5 ml of standard corn-meal fly medium and incubated at 28°C. To assess development of the larvae post-infection pupa were count every 12 hours for 5 days. For CFU counts, larvae were inoculated as described above. At each time point, 5 larvae were randomly collected, surface sterilized for 10 seconds in ethanol 70% and washed with miliQ water. Individual larvae were then transferred to Eppendorfs containing 300µl of 1x PBS and homogenized with a blender. The homogenate was diluted 100-fold and serial dilutions were plated in LB. Plates were incubated overnight at 30°C.

### Statistical analysis

Statistical analyses were performed in R(91) and graphs were generated using the package ggplot2(92) and GraphPad. All experiments were analysed using linear mixed-effect models [package lme4(93)]. Significance of interactions between factors was tested by comparing models fitting the data with and without the interactions using analysis of variance (ANOVA). Models were simplified when interactions were not significant. Multiple comparisons of the estimates from fitted models were performed with a Tukey HSD (honestly significant difference) test (packages lmerTest(94) and multicomp(95)). To each statistical group a letter is attributed, different letters stand for significant statistical difference.

## Data availability

Data will be fully available and without restriction upon request.

## Acknowledgments

We thank Joana Amaro for technical assistance, Rita Valente, Vitor Cabral, Roberto Balbontín, Tanja Dapa and André Carvalho for suggestions and helpful comments on the manuscript. We are very grateful to Bruno Lemaitre (EPFL) for sharing protocols and *Ecc15* strain.

## SUPPEMENTAL FIGURES

**Table S1.**
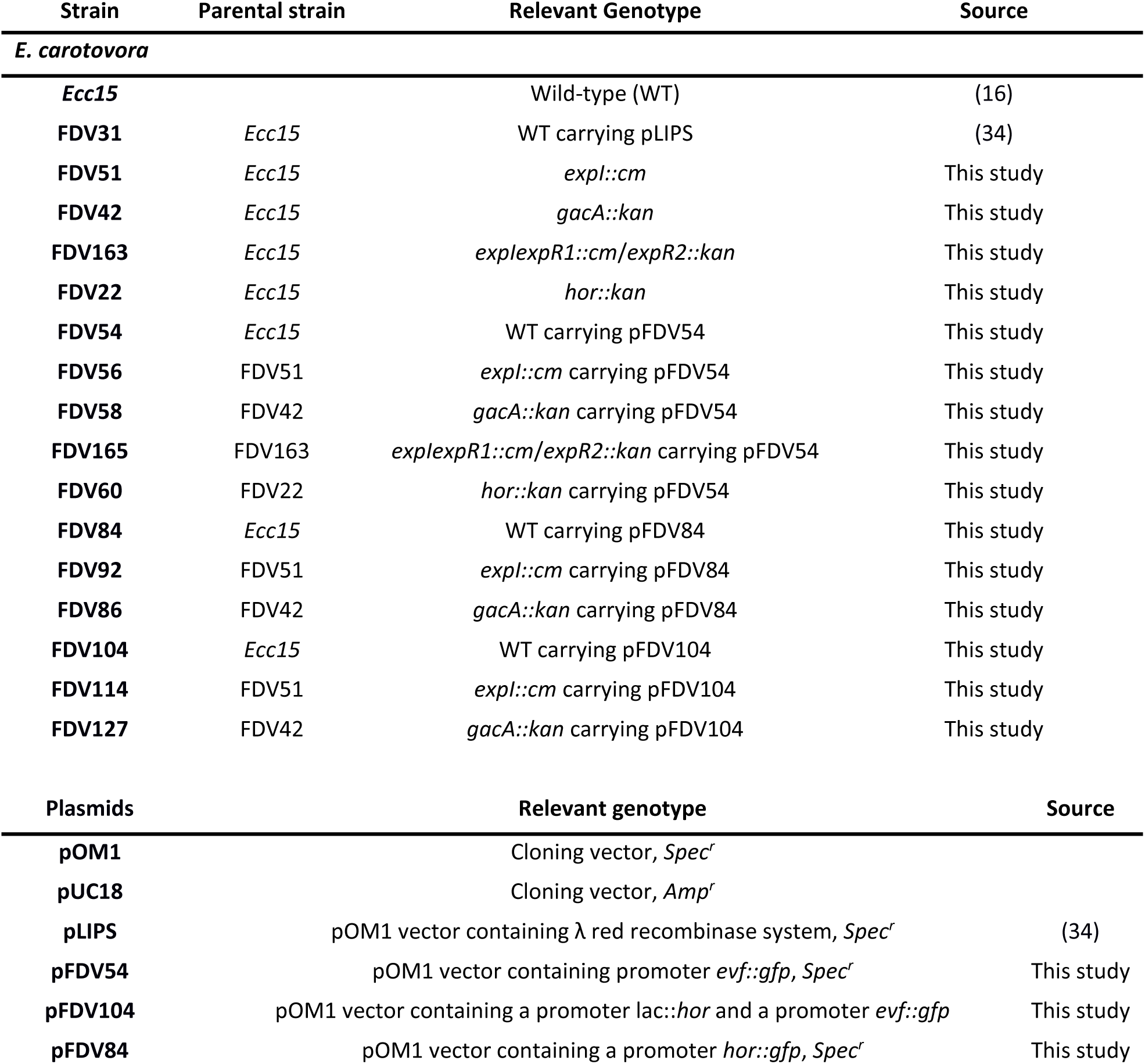
Strains and plasmids used in this study.

**Table S2.**
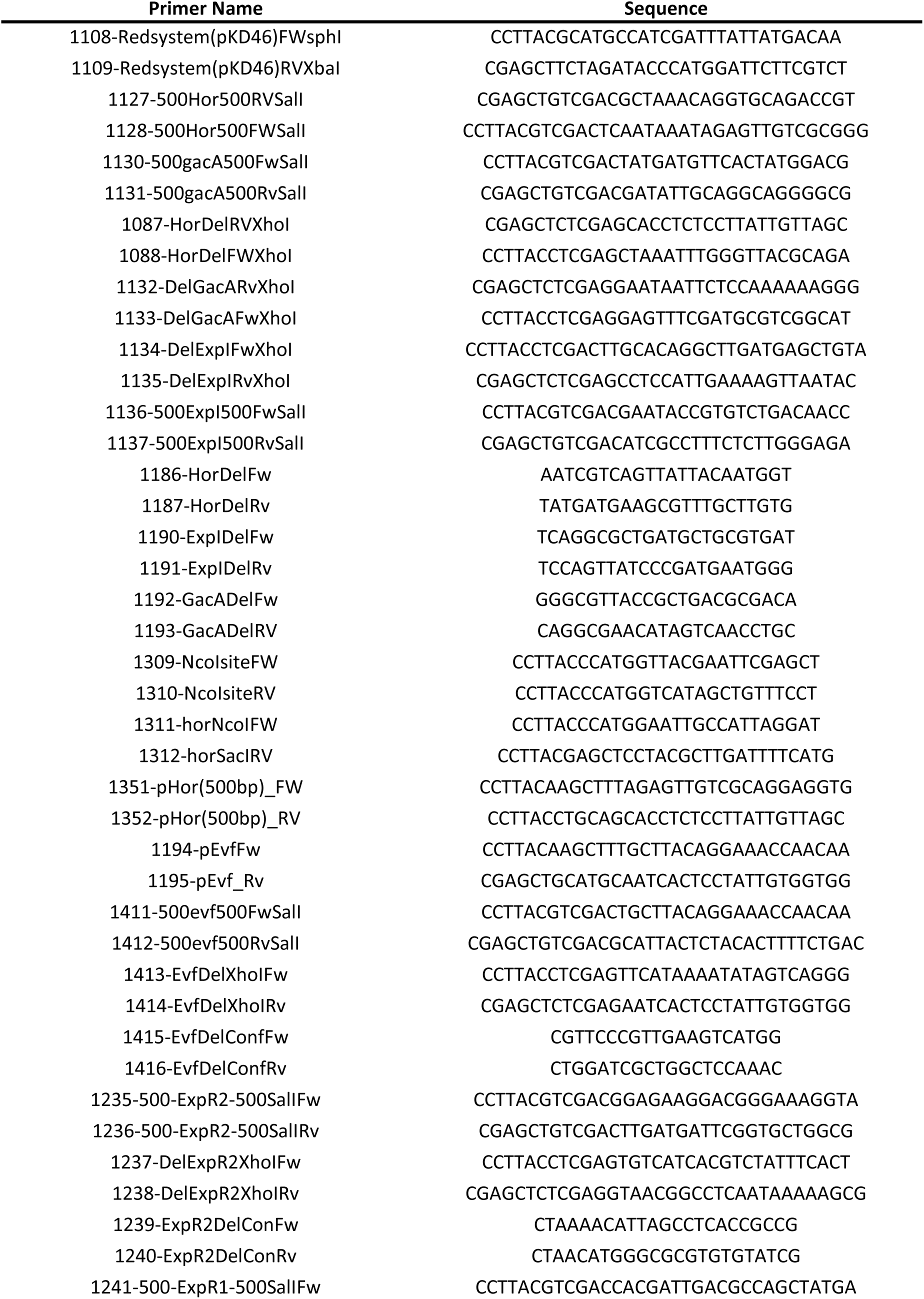
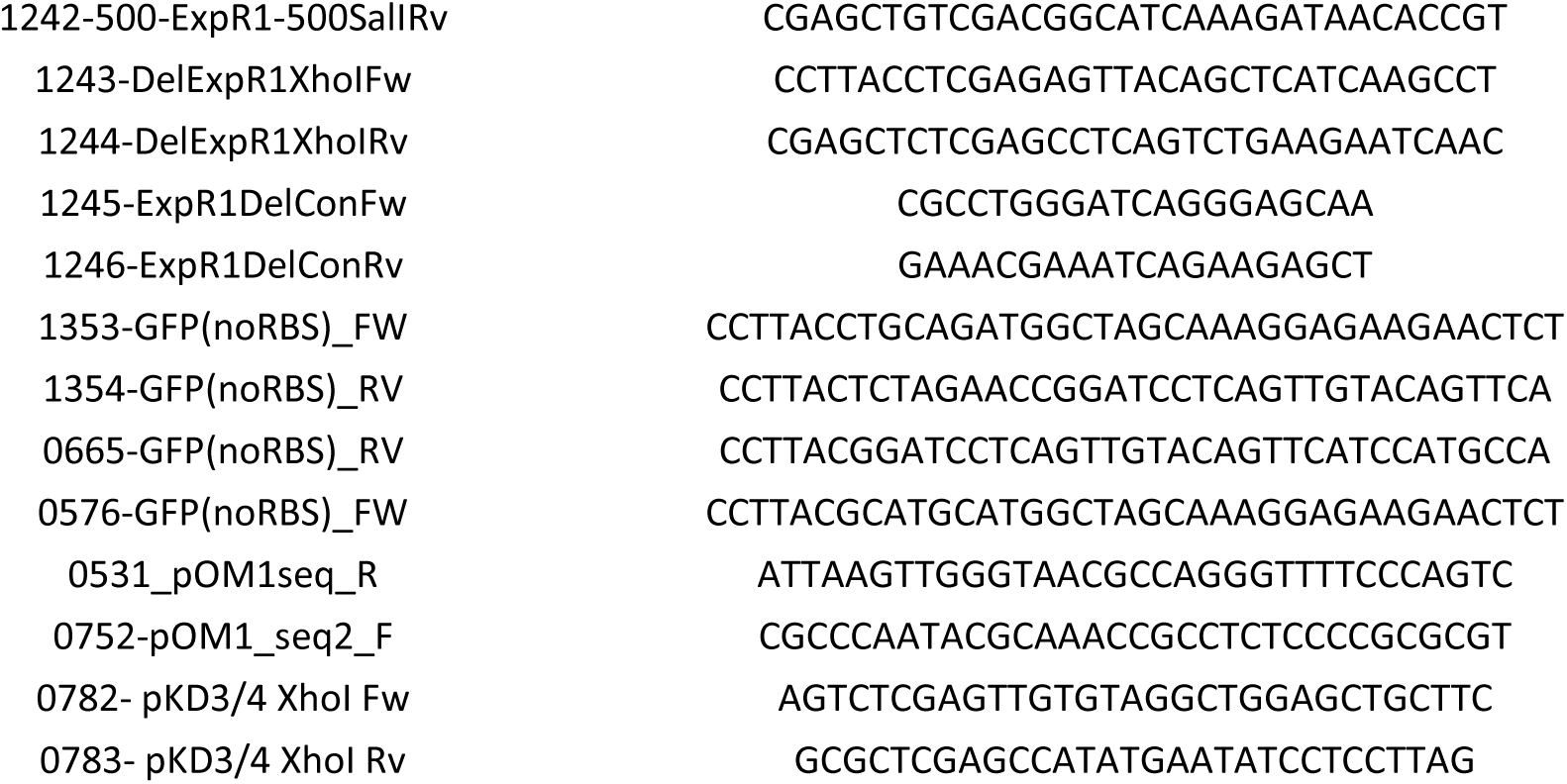
Primers used in this study.

**Table S3.** Orthologues of the Evf protein from *Erwinia carotovora Ecc15* present in the NCBI database (October 2019). The amino acid sequence from *Ecc15* was used as template to identify orthologues. All Proteins are defined as a complete match in the bidirectional best hits. Alignment template stands for the PDB sequence with the highest confidence used by phyre2 to predict orthologs secondary structure, corresponding to *Ecc15* Evf. All sequences were run in both phyre2 (72) and pfam database (96).

| Organism | Sequence ID | Locus Tag | Protein annotated function | % of amino acid query cover | % of amino acid identity | Alignment Template | % of confidence in the predicted structure |
| --- | --- | --- | --- | --- | --- | --- | --- |
| *P. carotovorum* Strain 14A | CP034276.1 | EIP93_02725 | Hypothetical protein | 100 | 100 | c2w3yB | 100 |
| *P. carotovorum* Strain Scc1 | CP021894.1 | SCC1_3840 | virulence factor | 100 | 100 | c2w3yB | 100 |
| *P. carotovorum* Strain 3-2 | CP024842.1 | OA04_05770 | virulence factor | 100 | 100 | c2w3yB | 100 |
| *Cedecea neteri* | WP_039302011 | LH86_RS13095 | Hypothetical protein | 100 | 70 | c2w3yB | 100 |
| *Enterobacter* sp. AG1 | WP_008453376 | A936_RS00125 | Hypothetical protein | 99 | 69 | c2w3yB | 100 |
| *Rahnella* sp. AA | WP_101079538 | CWS43_23475 | Hypothetical protein | 100 | 68 | c2w3yB | 100 |
| *Klebsiella aerogenes* | WP_087858097 | B9037_RS05845 | Hypothetical protein | 100 | 66 | c2w3yB | 100 |
| *Escherichia coli* | WP_113374258 | DUL12_RS15125 | Hypothetical protein | 100 | 66 | c2w3yB | 100 |

**Fig. S1.**
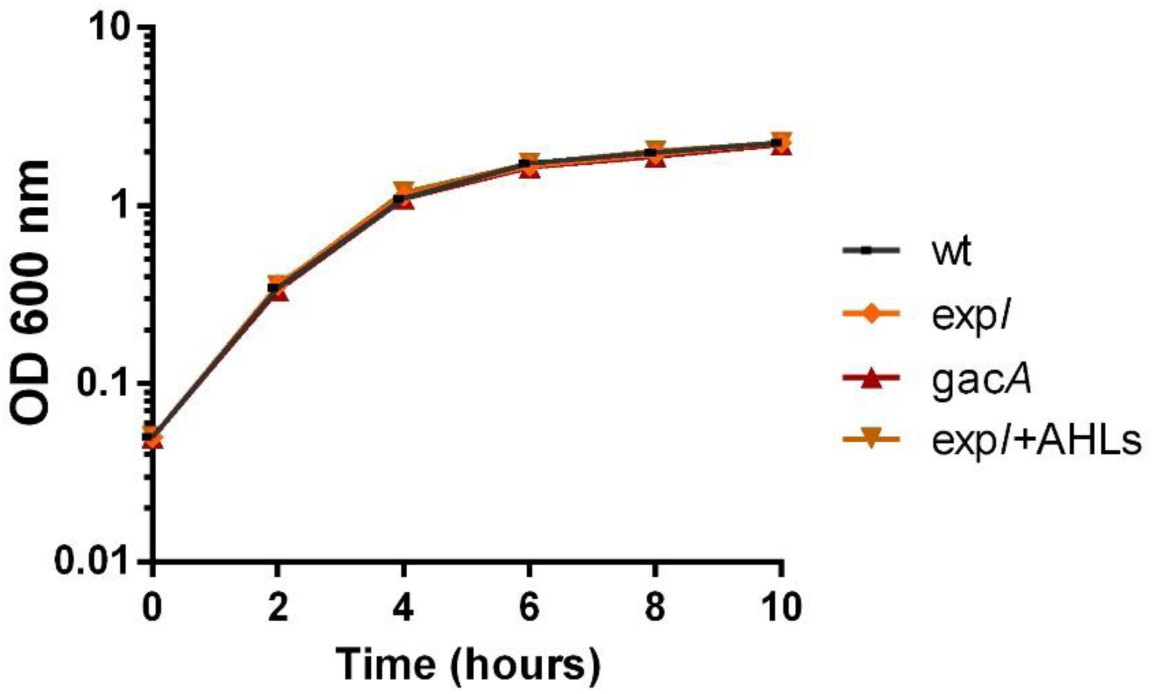
Growth curves of WT *Ecc15*, *expI* and *gacA* mutants carrying a P*_evf_*::*gfp* reporter fusion.

**Fig. S2.**
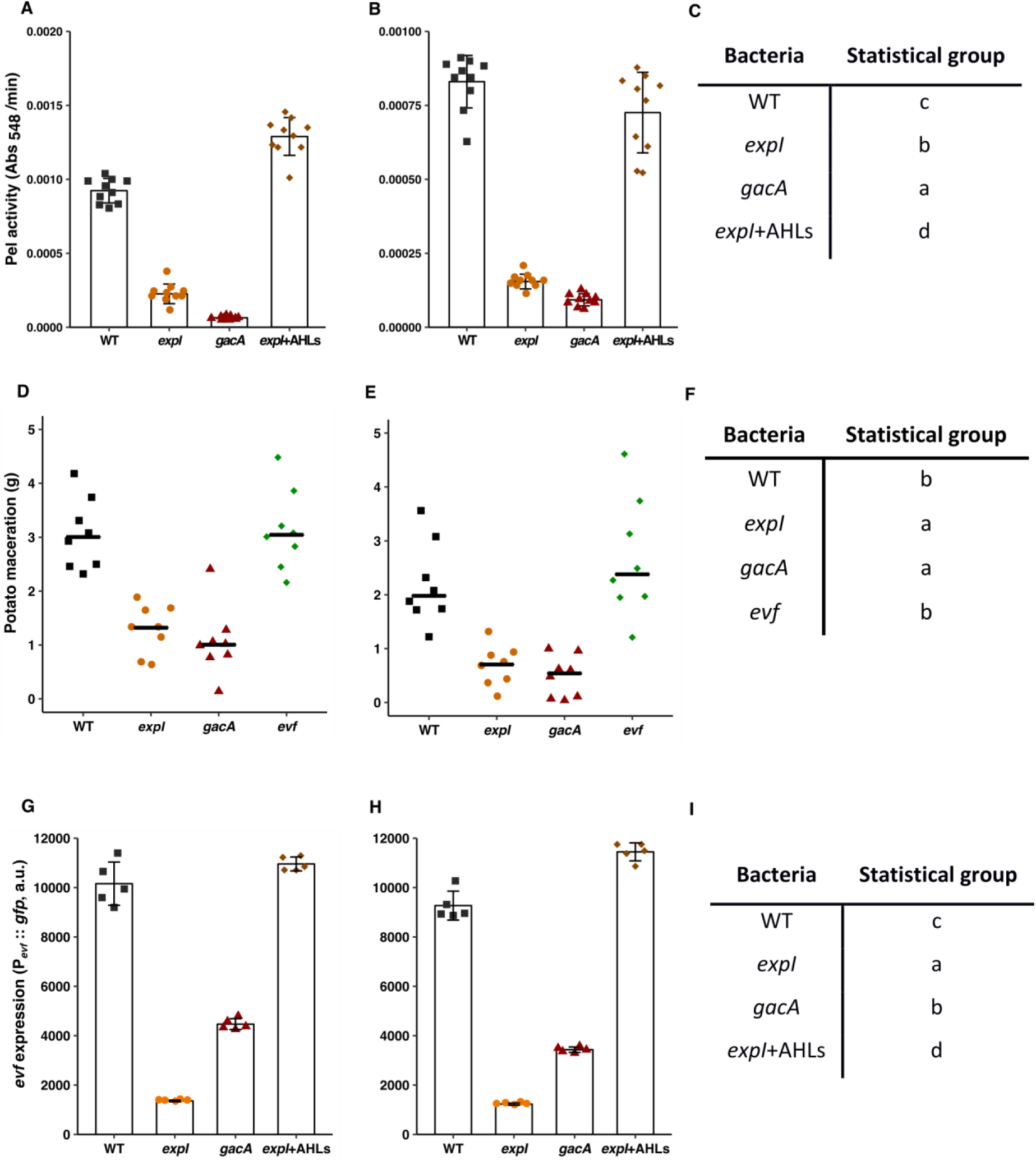
Independent replicates of the experiments shown in Fig. 1 (Production of pectate lyase and expression of *evf* is dependent on both quorum sensing and the GAC system). **(A, B)** replicates of experiments shown in Fig. 1A, **(C)** Statistical groups of all three experiments from Fig. 1A, **(D, E)** replicates of experiments shown in Fig. 1B, **(F)** Statistical groups of all three experiments from Fig. 1B, **(G, H)** replicates of experiments shown in Fig. 1C, **(I)** Statistical groups of all three experiments from Fig. 1C. Statistical analysis was performed using a linear mixed effect model. A Tukey HSD test was applied for multiple comparisons using the estimates obtain from the model.

**Fig. S3.**
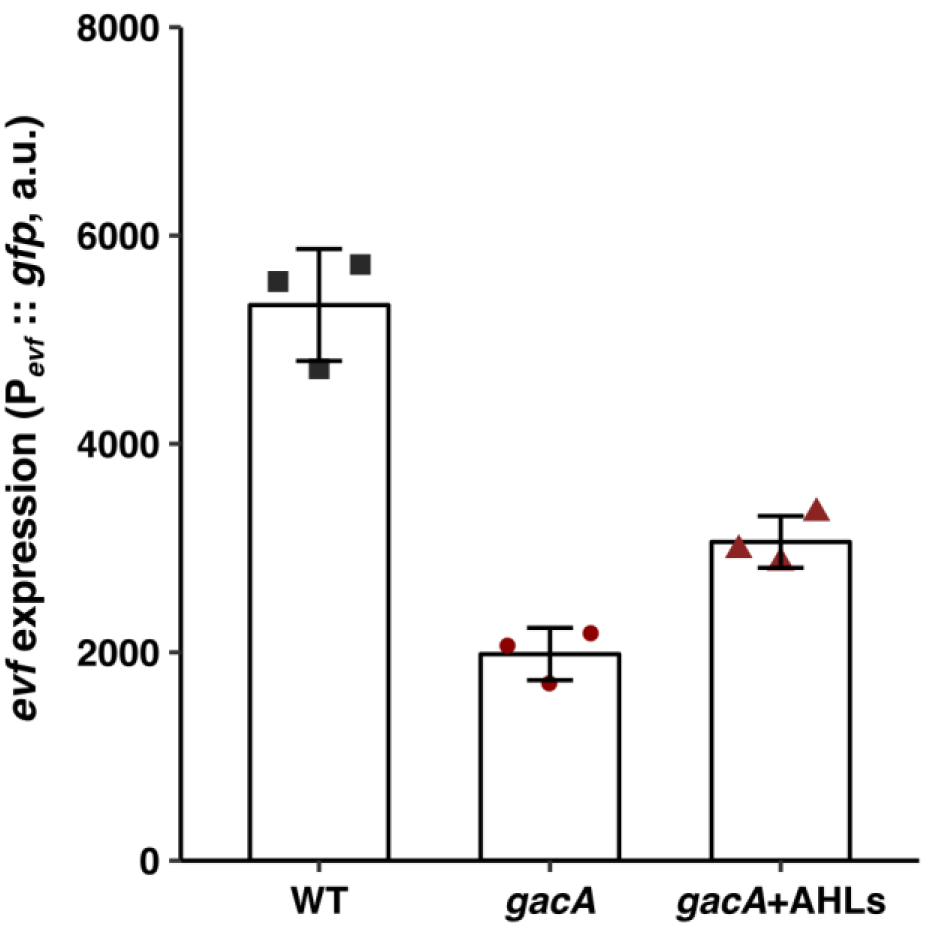
AHLs cannot complement intermediate levels of Pevf::gfp expression in a *gacA* mutant. P*_evf_*::*gfp* expression in WT *Ecc15* and *gacA* mutant at 6 hours of growth in LB + Spec. n=3. Complementation with AHLs was performed with a mixture of 1uM 3-oxo-C6-HSL and 3-oxo-C8-HSL. Error bars represent standard deviation of the mean.

**Fig. S4.**
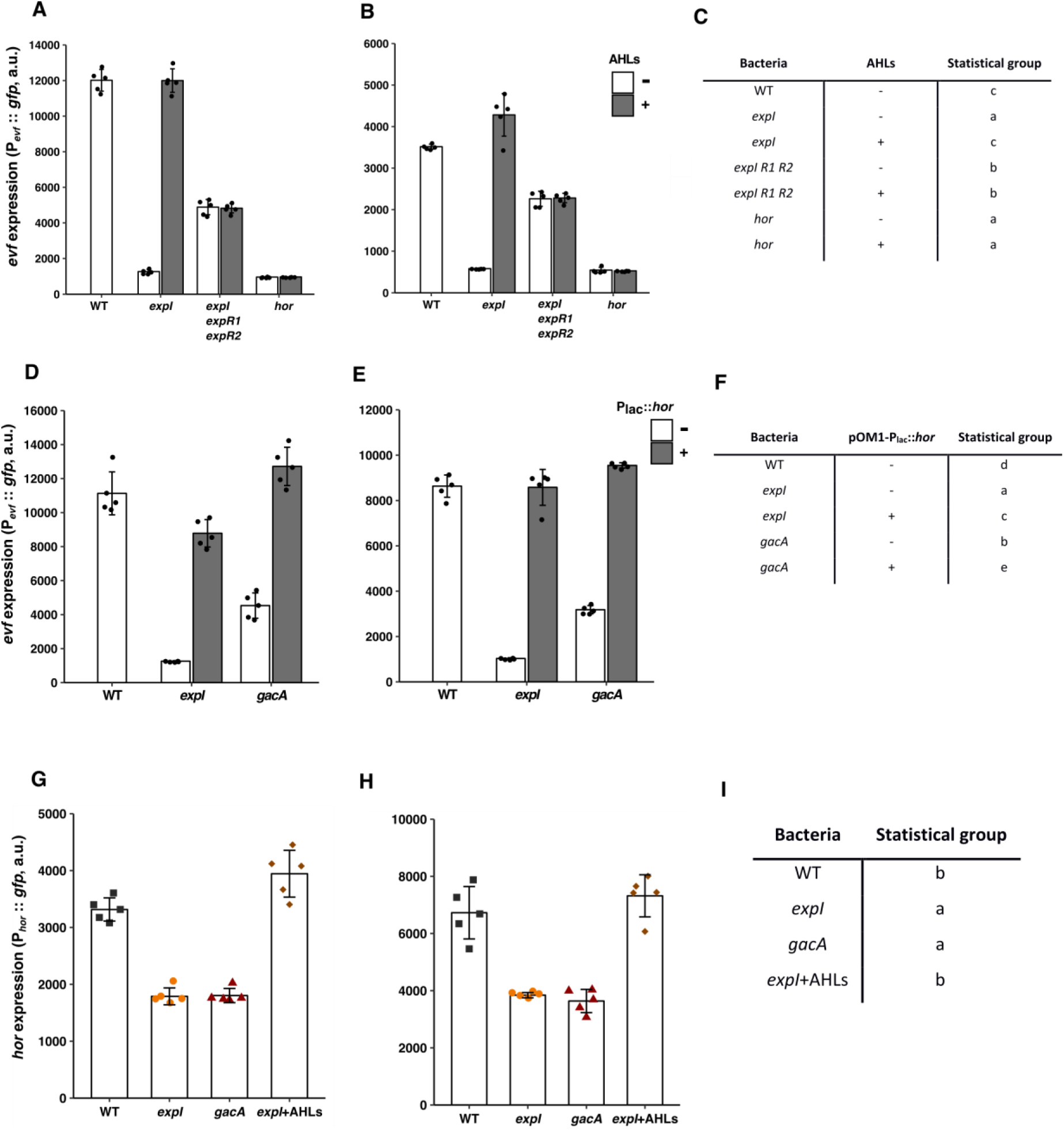
Independent replicates of the experiment shown in Fig. 2a (*evf* regulation by quorum sensing is dependent on ExpR receptors and *hor*). **(A, B)** replicates of experiments shown in Fig. 2A, **(C)** Statistical groups of all three experiments from Fig. 2A, **(D, E)** replicates of experiments shown in Fig. 2B, **(F)** Statistical groups of all three experiments from Fig. 2B, **(G, H)** replicates of experiments shown in Fig. 2C, **(I)** Statistical groups of all three experiments from Fig. 2C. Statistical analysis was performed using a linear mixed effect model. A Tukey HSD test was applied for multiple comparisons using the estimates obtain from the model.

**Fig. S5.**
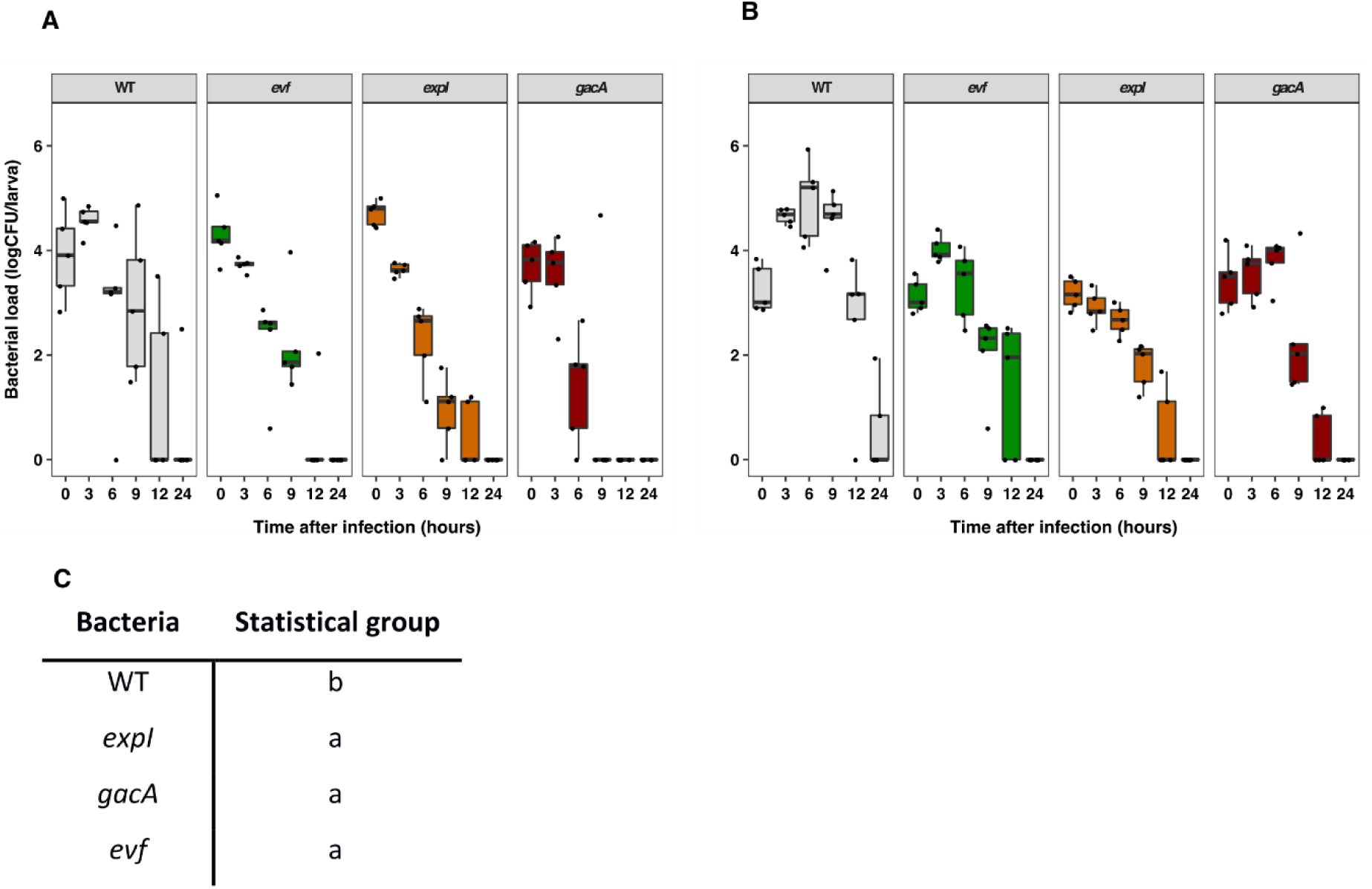
Independent replicates of the experiment shown in Fig. 3 (*Ecc15* loads are higher in *D. melanogaster* larvae orally infected with WT than with mutants impaired in *evf* expression.) **(A, B)** replicates of experiments shown in Fig. 3, **(C)** Statistical groups of all three experiments from Fig 3. Statistical analysis was performed using a linear mixed effect model. A Tukey HSD test was applied for multiple comparisons using the estimates obtain from the model.

**Fig. S6.**
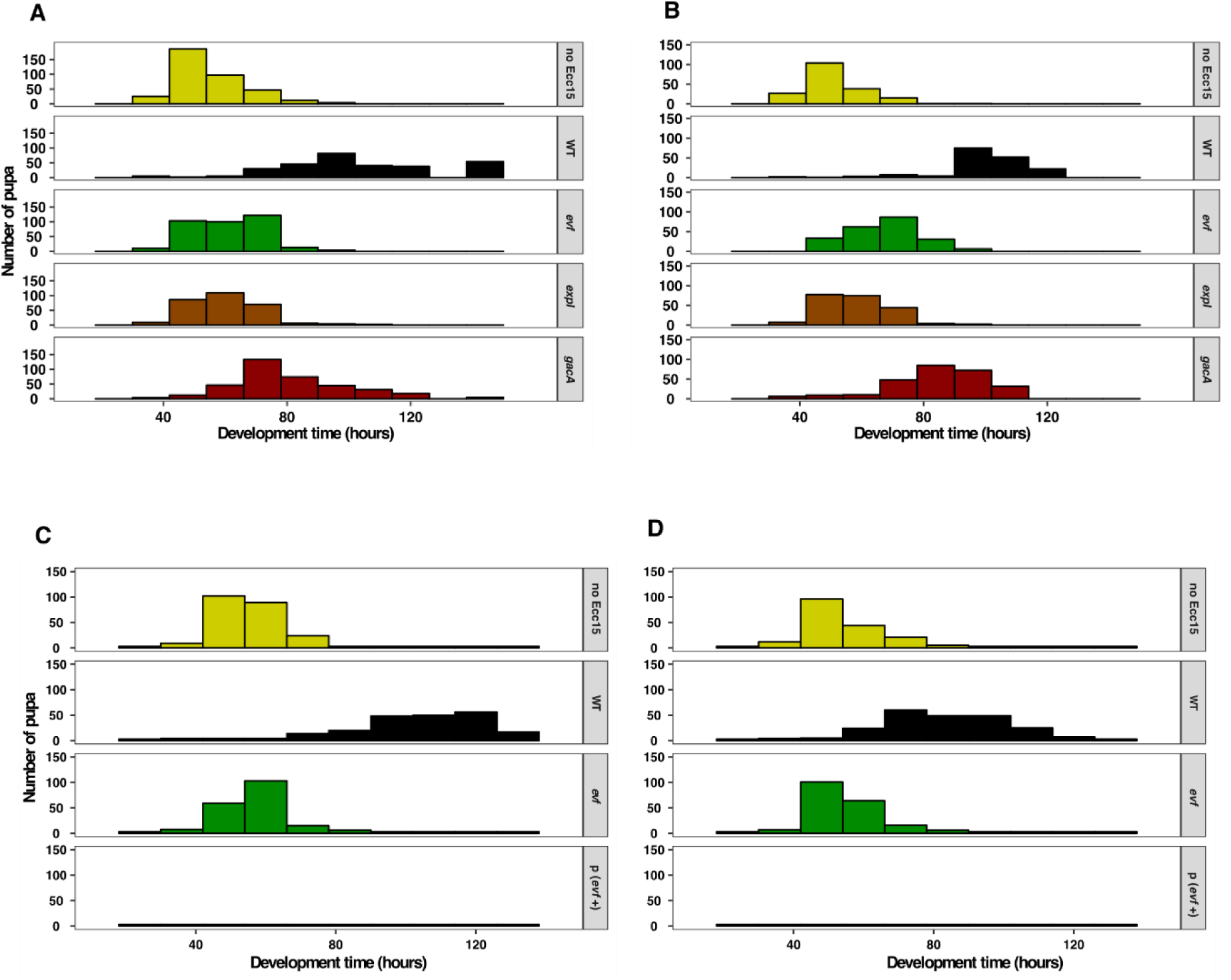
Independent replicates of the experiment shown in Fig. 4 (*Ecc15* causes a developmental delay in *D. melanogaster* larvae that is dependent on *evf*, quorum sensing and the GAC system). **(A, B)** replicates of experiments shown in Fig. 4A, **(C, D)** replicates of experiments shown in Fig. 4C.

